# EpiMolBio: A Novel User-Friendly Bioinformatic Program for Genetic Variability Analysis

**DOI:** 10.1101/2025.02.21.639439

**Authors:** Roberto Reinosa, Paloma Troyano-Hernáez, Ana Valadés-Alcaráz, África Holguín

## Abstract

**Motivation:** Genetic sequence analysis has become essential in many medicine, biology, and epidemiology fields. However, the currently available tools can pose a challenge for users without advanced computational skills.

**Results:** We present EpiMolBio, a free-to-use software designed with an intuitive, user-friendly interface that enables a broad spectrum of users to explore genetic variability. Its diverse toolkit encompasses sequence processing, conservation and variability analysis, consensus sequence generation, and genome mutation or amino acid changes identification, including specialized tools for HIV and SARS-CoV-2 analysis.

**Availability:** Freely available on the web at https://www.epimolbio.com and https://github.com/EpiMolBio/EpiMolBio.

**Supplementary information:** Supplementary Table 1, 2, and Supplementary Text 1.

## Introduction

Genetic sequence analysis plays a crucial role in the field of modern medicine, with heightened importance during the COVID-19 pandemic (Ray et al. 2021). The sequencing of microorganisms has become more common, with an increasing number of public databases (Elbe and Buckland-Merrett 2017, Los Alamos National Laboratory 2024). This fosters global research and collaboration to comprehend new strains and infectious agents, enabling variant identification, molecular evolution studies, and detection of conserved motifs, providing valuable information to develop diagnostic tools and vaccines, identify potential drug targets and drug resistance mutations (DRM).

However, many bioinformatics tools are costly or intricate, narrowing their user base and limiting global research collaboration. Our aim was to develop a user-friendly, versatile program for sequence analysis that can adapt to the user’s objectives. This program was initially developed in 2018 for HIV variability and drug-resistance studies (Troyano-Hernáez, Reinosa and África Holguín 2022, Troyano-Hernáez et al. 2021, Troyano-Hernáez, Reinosa and Africa Holguín 2022, Valadés-Alcaraz, Reinosa and Holguín 2022). At the beginning of the COVID-19 pandemic, new functions were developed for SARS-CoV-2 mutation tracing and variant analysis (Troyano-Hernáez, Reinosa and Holguín 2021, Troyano-Hernáez, Reinosa and África Holguín 2022), together with supplementary software for massive alignments, fasta file editing, and sequence processing (Valadés-Alcaraz et al. 2024).

Although initially developed and tested for viral genome analysis (HIV, SARS-CoV-2), the software supports sequences in the widely-used .fasta format, along with user-defined reference sequences, enabling the study of a wide range of pathogenic organisms and other significant genomic entities in medical and biological research. These capabilities allow researchers to conduct precise genomic analyses tailored to their specific needs, significantly advancing our understanding and treatment of various diseases. To date, we have utilized this software in collaboration with other scientists across various research projects. These include designing and optimizing primers for HCV PCR-based amplifications, developing aptamers for the molecular diagnosis of SARS-CoV-2, RSV and Clostridium difficile, among other pathogens, studying the conservation of C. difficile toxins and viral proteins (HIV, RSV, SARS-CoV-2), among other applications.

## Methods

EpiMolBio is developed in the Java programming language using Apache NetBeans 14 as the Integrated Development Environment (IDE). The graphical interface has been developed using Java Swing. The ‘Multiple alignments’ function uses the MUSCLE program (Edgar 2004) for performing alignments. Additionally, the ‘Mutation frequency’ and ‘Partial similarity’ functions employ the BioJava library for alignments. Apache Maven 3.8.5 is used for project management and build automation.

The program is freely available at epimolbio.com and is registered under the Creative Commons Attribution-NonCommercial-NoDerivatives 4.0 license (number 2305114294344). The source code for EpiMolBio is available on GitHub at https://github.com/EpiMolBio/EpiMolBio, with brief comments provided in both English and Spanish to facilitate understanding. The software executable and additional documentation can be accessed on the official website at www.epimolbio.com. The website, available in both Spanish and English, includes links to a complete user manual in both languages. EpiMolBio is a portable software that only requires the prior installation of Java 11 (Java | Oracle) to function.

## Results

EpiMolBio is a free bioinformatics program for the analysis of genetic and protein sequences that can be used for structural, functional or evolution studies. It has been designed to be user-friendly, with powerful tools and customizable workflows. The EpiMolBio interface is designed to be as simple and intuitive as possible, eliminating the need for prior programming knowledge or advanced bioinformatics skills (**Figure 1**).

**Figure 1.**
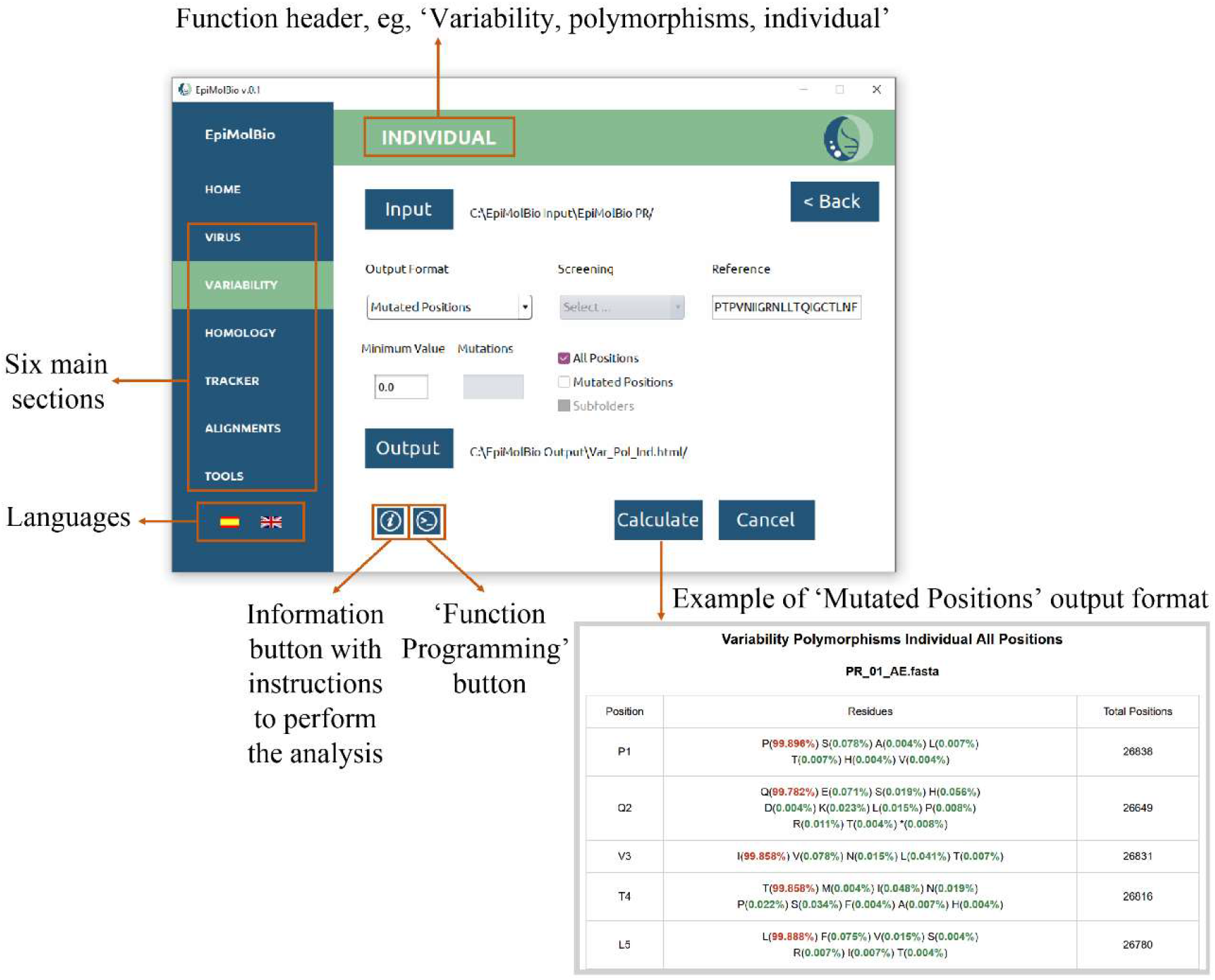
Example of EpiMolBio interface.

The software is available in two languages: English and Spanish. Each function features an information button that provides a summary of the steps required to carry out the analysis and details on the supported output formats. The website features a comprehensive user manual in both languages, offering detailed information about each function and step-by-step instructions for all tasks. Users can also access example datasets and expected outputs for EpiMolBio’s various functions for training purposes in the following GitHub link (https://github.com/EpiMolBio/EpiMolBio) and webpage link (https://epimolbio.com/manual-en/), allowing them to download sample data and generate different types of results (**Supplementary Table 1**). Additionally, a contact form is available for assistance, and video tutorials will be uploaded soon.

To input data, users must select the folder containing the .fasta sequences. Each .fasta file can include one or more sequences, allowing for the simultaneous analysis of multiple sequences. An exception to this occurs during subsequent rounds of the ‘Consensus’ tool, which generates consensuses of consensuses. In these rounds, the outputs from the previous round, saved in .txt format, are used as input. Rather than selecting individual files, users must choose the folder containing the relevant .fasta or .txt files. It is essential that the folder contains only the files to be analyzed, as individual files are not displayed. Failure to adhere to this requirement may cause errors or improper processing.

Additionally, the program supports the input of folders containing subfolders for specific functions. This allows for the joint analysis of multiple .fasta files organized by variants, date, or other criteria. By handling complex folder structures, the software becomes more flexible and efficient for genomic sequence analysis. Simply select the main folder containing the subfolders as the input.

EpiMolBio provides a range of output formats, often offering multiple output options for the same function, allowing users to customize results. Outputs are in .fasta format for sequence modi-fication and localization functions, and .html or .csv tables for analytical results, except for ‘Consensus’, which is a .txt file. The .html output format can be viewed in a web browser and copied into Excel or Word for modifications. Results in .csv format can be easily edited in Excel or similar programs to suit the user pref-erences. In several output files, residues, percentages, or table cells will be color-coded based on their frequency percentage, making it easier to identify the polymorphisms occurrence at a glance. Epi-MolBio also enables workflow automation to streamline tasks and save time.

The program is organized into six distinct sections, each containing several functions (**Figure 2**). The detailed description of each function is available in the **Supplementary Table 2**, an illustrative example of how EpiMolBio is used is provided in **Supplementary Text 1**.

**Figure 2.**
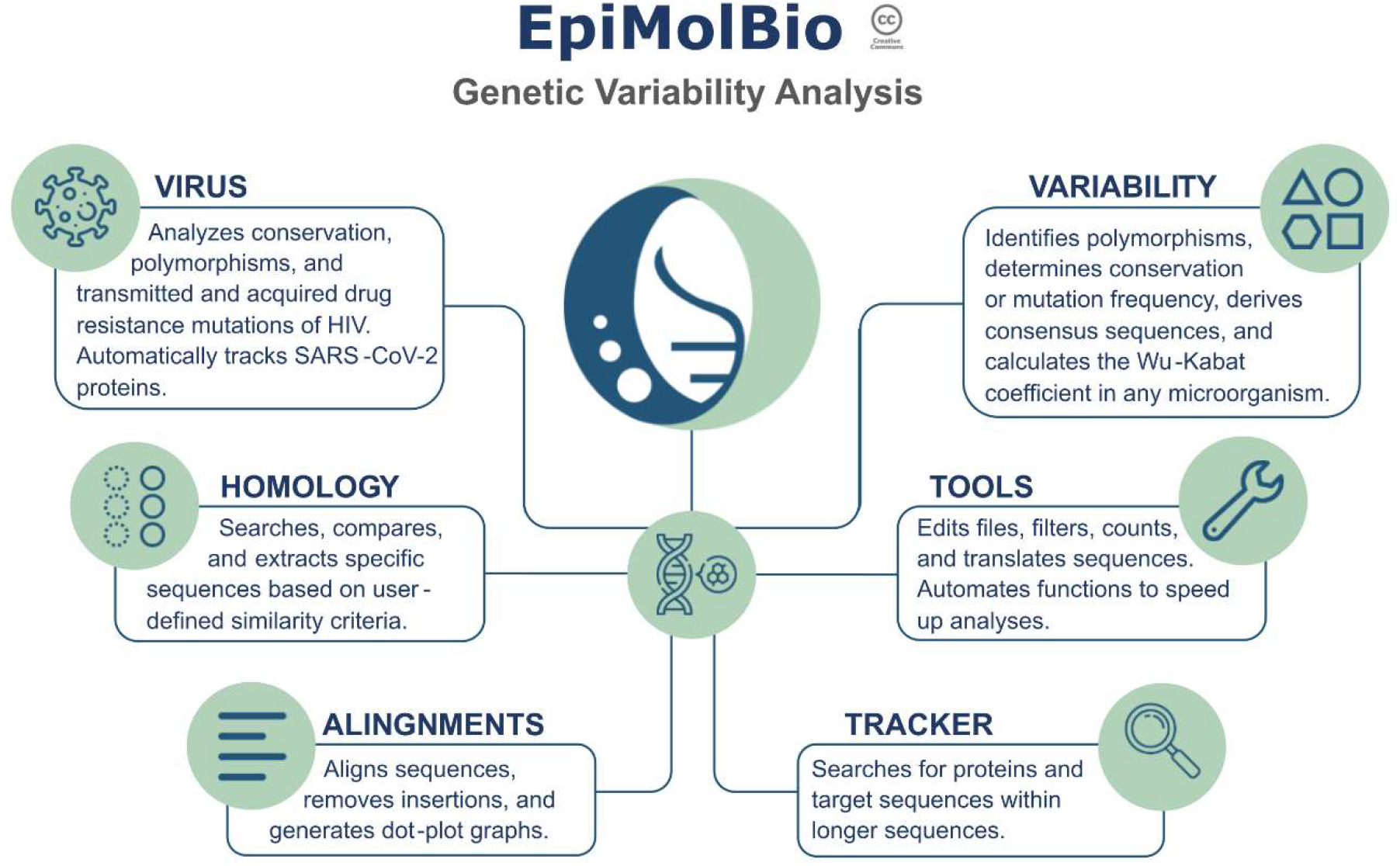
Summary of the six EpiMolBio main sections.

### Virus section

The Virus section specializes in HIV and SARS-CoV-2 analysis (**Table 1**). The HIV ‘Resistance Mutations’ function detects and calculates the frequency of acquired DRM in HIV-1 through individual and codon-based assessments, using the Stanford HIV Drug Resistance Database v9.7 updated in November 9th 2024, DRM list (Stanford University 2024), and following its DRM classification (Stanford University 2024). It also analyzes HIV-1 transmitted DRM, based on the WHO list of mutations for protease and reverse transcriptase (Bennett et al. 2009) and Tzou et al paper for integrase (Tzou, Rhee, et al. 2020). For HIV-2, DRM analysis is based on the HIV-2 EU Tool v.2 (Charpentier et al. 2015) and recent literature sources(Stanford University 2024, Tzou, Descamps, et al. 2020). The DRM lists within the program will be periodically updated.

**Table 1.** Summary of EpiMolBio Virus section functions.

| VIRUS |  |
| --- | --- |
| HIV, Resistance Mutations | Calculates the percentage of acquired/transmitted DRM of HIV Pol proteins and p24 (Capsid protein). |
| HIV, Other Pol Mutations | Detects any mutation (not only DRM) in HIV-1 or HIV-2 Pol sequences and their frequency. |
| HIV, Pol Conservation | Displays the most prevalent AA and its percentage for each position of a selected Pol protein sequence. |
| SARS-CoV-2 Protein Tracker | Retrieves the sequences of one or many selected SARS-CoV-2 proteins from complete genomes. |

The HIV ‘Other Pol Mutations’ function detects any mutations within the pol gene of HIV-1 and HIV-2, not limited to DRMs. The ‘Pol Conservation’ function provides tools for analyzing the conservation of HIV Pol proteins, generating a consensus sequence from input sequences and identifying the most conserved residue for each position along with its frequency with a color code. Additionally, the ‘SARS-CoV-2 Protein Tracker’ allows input of complete genomes to retrieve sequences for each viral protein in either amino acid or nucleotide format.

### Variability section

This section offers various specialized tools for analyzing genetic sequence variability across diverse organisms, allowing users to input their own reference sequence (**Table 2**). The ‘Polymorphisms’ function enables the detection of genetic polymorphisms, providing their location and frequency through individual and codon-based assessments. It includes tools for detecting, pinpointing, and quantifying mutations at specific or multiple positions. Users can define specific mutations of interest and set a minimum occurrence frequency for displaying results. Additionally, users can identify mutations unique to individual input files using the ‘Markers’ tool to discover variant-specific mutations or analyze mutation combinations with the ‘Multiple Mutations’ tool.

**Table 2.** Summary of EpiMolBio Variability section functions.

| VARIABILITY |  |
| --- | --- |
| Polymorphisms* | **Mutated Positions** and **Mutations Table** detect and locate polymorphisms, and their frequency using any sequence as a reference.<br>**Markers** detects mutations exclusive to each file compared to the rest of the input files.<br>**Multiple Mutations** detects mutation combinations providing their frequency.<br>**Codon function** detects codons different from those in the reference sequence and their frequency.<br>**Mutations by Position** detects the residues at one position or several combined positions providing their frequency. |
| Conservation* | Reports the most prevalent residues and their frequency. Additionally, it generates consensus sequences.<br>**Codon function** presents the most conserved codon from a nucleotide sequence. |
| Consensus | Provides consensus sequences and consensus of consensus performing multiple rounds of analysis. |
| Wu-Kabat Coefficient | Provides the Wu-Kabat variability coefficient of protein sequences to study the susceptibility of an AA position to evolutionary replacements. |
| Mutation Frequency | Generates a set of parameters related to the frequency of mutations in a group of sequences. |

The ‘Conservation’ function evaluates sequence conservation by identifying the most common residue or codon and its percentage of occurrence. It also allows users to apply filters to refine frequency parameters. The ‘Consensus’ function enables users to derive consensus sequences from .fasta input files, which may include sequences generated from prior consensus analyses. Multiple rounds of successive analysis can be conducted to derive consensuses of consensuses.

The ‘Wu-Kabat Coefficient’ function enables the study of amino acid position susceptibility to evolutionary replacements using the Wu-Kabat formula. Finally, the ‘Mutation Frequency’ function computes parameters related to mutation frequency in a group of nucleotide or amino acid sequences.

### Homology section

This section facilitates sequence comparison, peptide detection, and identification of conserved motifs, useful for designing primers, diagnostic and therapeutic aptamers, and vaccines (**Table 3**). It includes three tools. ‘Similarity’ searches for a specific short sequence entered by the user within the input file sequences, and determines its frequency. ‘Partial Similarity’ compares a user-entered sequence with sequences from the input file, identifying similar regions based on a defined percentage of similarity. ‘Search for Conserved Sequences’ extracts conserved sequence fragments from a set of input sequences, allowing users to search within a specific region, choose the fragment length, and define the conservation percentage.

**Table 3.**
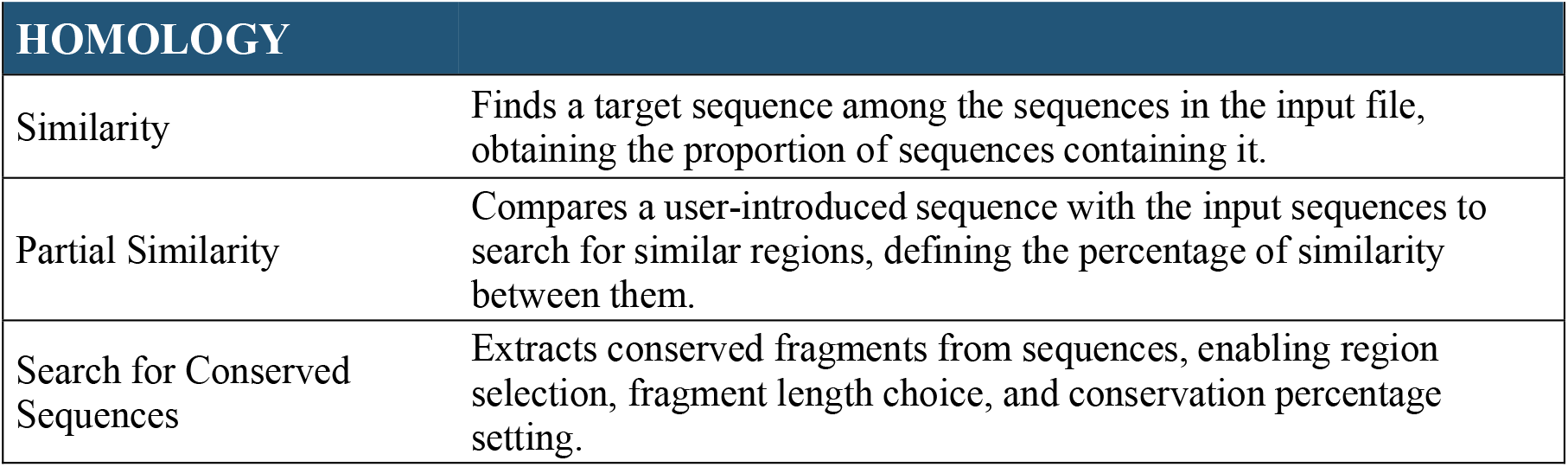
Summary of EpiMolBio Homology section functions.

| HOMOLOGY |  |
| --- | --- |
| Similarity | Finds a target sequence among the sequences in the input file, obtaining the proportion of sequences containing it. |
| Partial Similarity | Compares a user-introduced sequence with the input sequences to search for similar regions, defining the percentage of similarity between them. |
| Search for Conserved Sequences | Extracts conserved fragments from sequences, enabling region selection, fragment length choice, and conservation percentage setting. |

### Tracker section

The ‘Tracker’ section locates sequence fragments within longer sequences, enabling the identification of specific proteins or genes within a genome (**Table 4**). Users can search for fragments based on similarity to a user-entered reference sequence or using flanking sequences to locate proteins.

**Table 4.**
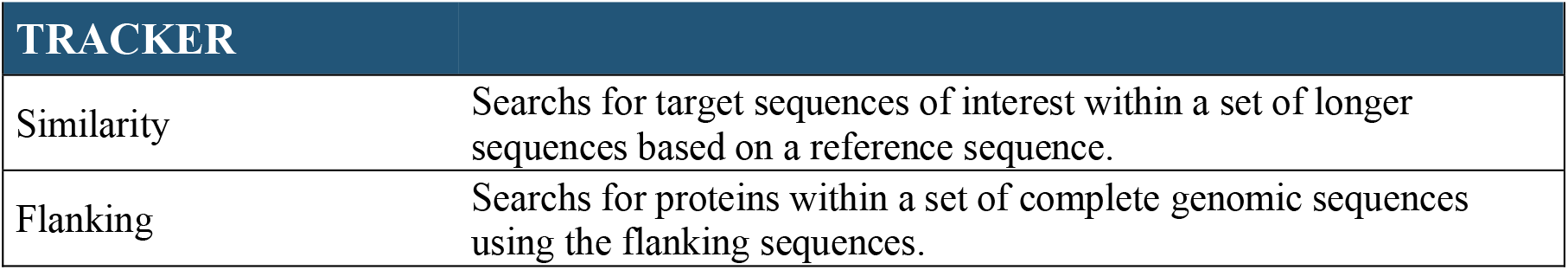
Summary of EpiMolBio Tracker section functions.

### Alignments Section

This section allows users to perform multiple alignments using the MUSCLE v3.8.31 program, generate dot plot charts, and remove insertions (**Table 5**). EpiMolBio offers customizable multiple alignment options tailored to the input file type. Users can choose to align sequences in batches or individually (pairwise alignment), depending on factors such as sequence length, quality, and expected mutation count, which speeds up the process and enables the alignment of thousands of sequences. It is important to note that when sequences are individually aligned against a reference, the process follows a pairwise alignment scheme. However, in the final output file, all sequences are aligned with each other after the removal of insertions. This occurs because, when aligning each sequence with the reference, any insertion present in the sequence introduces gaps in the reference. These gaps serve as guides to identify and eliminate the insertions in the aligned sequence. By repeating this procedure with all sequences, the final result is a set of sequences aligned with one another at a uniform length, facilitating their comparison and analysis.

**Table 5.**
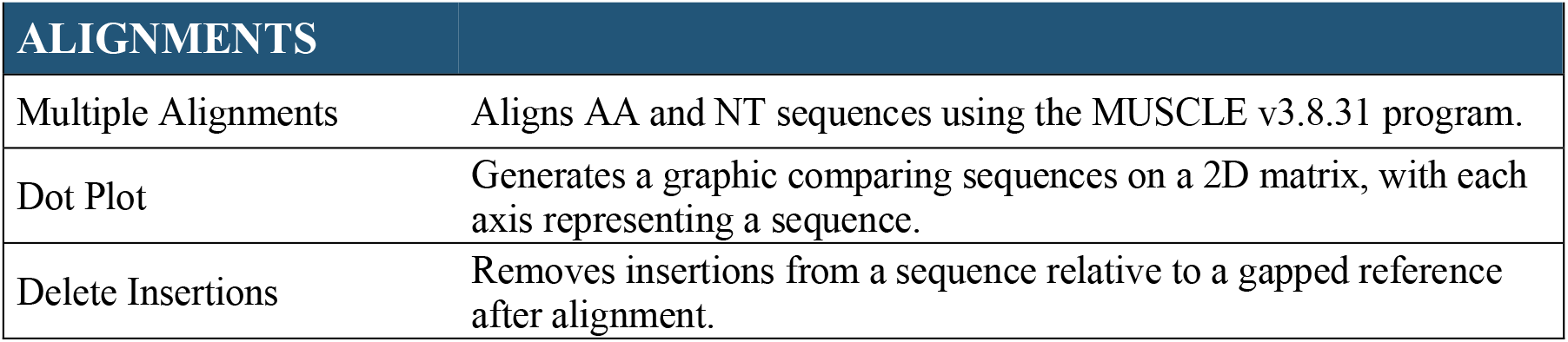
Summary of EpiMolBio Alignments section functions.

| ALIGNMENTS |  |
| --- | --- |
| Multiple Alignments | Aligns AA and NT sequences using the MUSCLE v3.8.31 program. |
| Dot Plot | Generates a graphic comparing sequences on a 2D matrix, with each axis representing a sequence. |
| Delete Insertions | Removes insertions from a sequence relative to a gapped reference after alignment. |

### Tools Section

This section offers a range of tools for file and sequence editing tasks, such as file merging, duplicate sequence removal, sequence filtering based on specific mutations, and character replacement in headers or genetic sequences (**Table 6**). Additionally, the ‘Filters’ tool enables sequence filtering based on quality and offers the ability to filter files by header parameters, separating them into different files according to the chosen parameter, and to filter sequences that have a specific set of characters in their headers, making it easy to classify sequences by date, country, or other criteria.

**Table 6.** Summary of EpiMolBio Tools section functions.

| TOOLS |  |
| --- | --- |
| File Editing |  |
| Merge Files | Combines multiple .fasta files into a single .fasta file. |
| Unique Sequences | Removes duplicated sequences from one or multiple input .fasta files. |
| Sequence Search | Filters sequences from .fasta files that contain one or multiple mutations chosen by the user. |
| Find and Replace | Replaces a series of characters in the header or the genetic sequence of one or multiple .fasta files. |
| Filters |  |
| Header Filtering | Filters .fasta files based on header parameters. |
| Specific Filter | Filters .fasta sequences based on specific characters in their headers. |
| Partial Sequence Filtering | Filters .fasta sequences by quality, based on "?" in AA or "N" in NT sequences. |
| Translation | Translates .fasta sequences from NT to AA. |
| Count Sequences | Counts total sequences in one or multiple .fasta files and identifies those with mutations. |
| Function Programming | Automates program functions by chaining them for sequential execution without manual input. |

Furthermore, this section offers sequence translation and counting functionalities, along with the ‘Function Programming’ tool, which automates and customizes workflows through sequential task execution with a CSV instruction file, resulting in significant time savings. This CSV file can be easily generated by clicking the ‘Function Programming’ button within each function’s interface.

## Conclusions

EpiMolBio is a user-friendly software designed to facilitate genetic sequence analysis of large datasets, initially tailored for viral genomes but adaptable to a wide range of microorganisms. The capability to input user-defined reference sequences enables the study of diverse microorganisms or other genomic entities with biological or biomedical interest. It features specific applications for HIV DRM identification and resistance data presentation, as well as SARS-CoV-2 proteins tracking. Alongside these features, EpiMolBio offers a comprehensive suite of tools for sequence alignment, editing, and workflow automation, simplifying complex analytical processes. Ongoing efforts are focused on expanding EpiMolBio’s utility to analyze other genomes of biological or medical relevance, including bacterial and human genes, to contribute to disease research.

## Acknowledgements

We thank the IRYCIS Innovation Unit for the help during the EpiMolBio registration and Val Fernández-Lanza, head of the Bioinformatic Unit at IRYCIS, for his helpful comments during the revision of the manuscript writing. We thank Fundación Familia Alonso for their collaboration in providing financial support for the development of the program.

## Funding

This work was partially supported by Fundación Familia Alonso [grant 2022/0027]; and by Fundación Ramón y Cajal para la Investigación sanitaria (FIBioHRC) [FONDOS FUR 2020/0285], plus fundrising activities and donations. P.T. was funded by Fundación Familia Alonso [grant 2022/0027]. R.R. was funded by Fundación Ramón y Cajal para la Investigación Sanitaria (FIBioHRC) [FONDOS FUR 2020/0285], and by Fundación Familia Alonso [grant 2022/0027].

This study is included in the Spanish Network CIBERESP (Centro de Investigación Biomedica en Red de Epidemiología y Salud Pública CB06/02/0053).

