## Supplementary Text 1 for "EpiMolBio: A Novel User-Friendly Bioinformatic Program for Genetic Variability Analysis"

Example of the use of the EpiMolBio program to identify 100% conserved fragments of 20-25 amino acids from a specific subunit of a viral protein:

In this example the aim was to identify 100% conserved sequence fragments of 20-25 residues within SARS-CoV-2 Spike protein subunit 1 (S1), region encompassing amino acids 14-685 of that protein. These could be used later for designing aptamers intended for COVID-19 treatment or diagnosis.

In the first place, we downloaded recent sequences from GISAID database (Shu and McCauley 2017). We retrieved 65 complete genomes available on GISAID from November 1st 2023 to November 7th 2023, via https://www.epicov.org/epi3/frontend#4201de.

Next, we used the 'SARS-CoV-2 Protein Tracker' function of EpiMolBio program to obtain the Spike proteins in amino acids from the 65 GISAID complete genomes. The input was the folder containing the 65 complete nucleotide sequences. Then, we selected the Spike protein and chose 'Translate' in the program interface. As output we created a new folder named 'Spike'. After the process, this folder contained 64 SARS-CoV-2 Spike proteins in .fasta format, since one sequence was lost due to being highly mutated. When an input sequence contains many mutations or is incomplete in the region that corresponds to the searched protein, the Protein Tracker cannot recognize it and therefore, the program will not retrieve it.

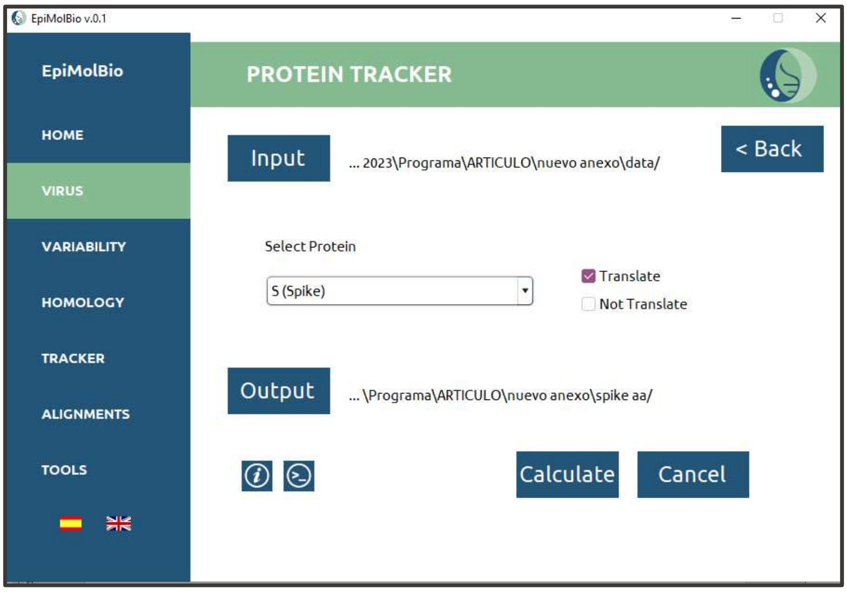


The next step was to align the sequences using the 'Multiple Alignments' function. Here, we used the previously created folder named 'Spike' as input folder. We chose 1 sequence per file, without preserving insertions and input the Spikes reference sequence (accession number NC_045512.2) without spaces or line breaks. For the output we created a new folder named 'S aligned'. After the process, this folder contained the 64 aligned Spike amino acid sequences.

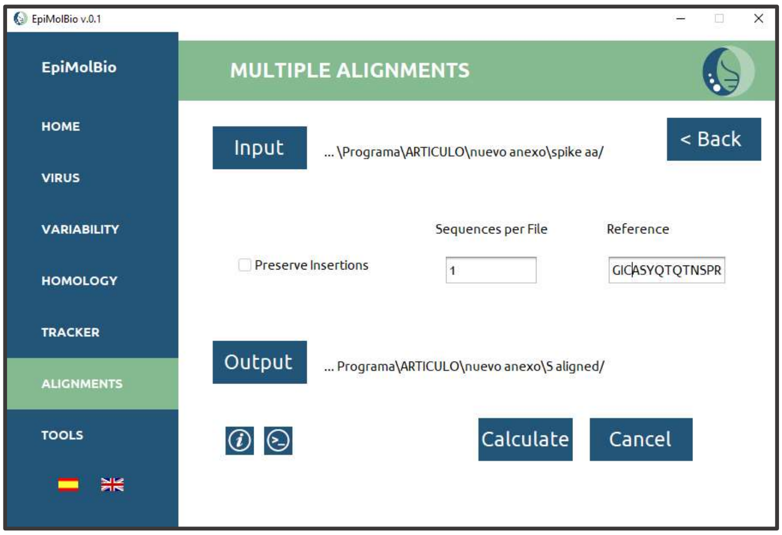


We wanted the Spike protein sequences to be complete. For this purpose, we used EpiMolBio 'Partial Sequence Filtering' in 'Tools'. This function allowed us to filter the sequences with unknown residues within a group of sequences. The input was the previous output folder. In the program's interface, we chose amino acids as the sequences had already been translated and typed 100.0 in 'Filter' as we only wanted to keep the complete sequences. Then, we created a new folder as output named 'Filtered'. We were left with 41 complete Spike sequences and an .html file showing how many sequences we had lost in the filtering process.

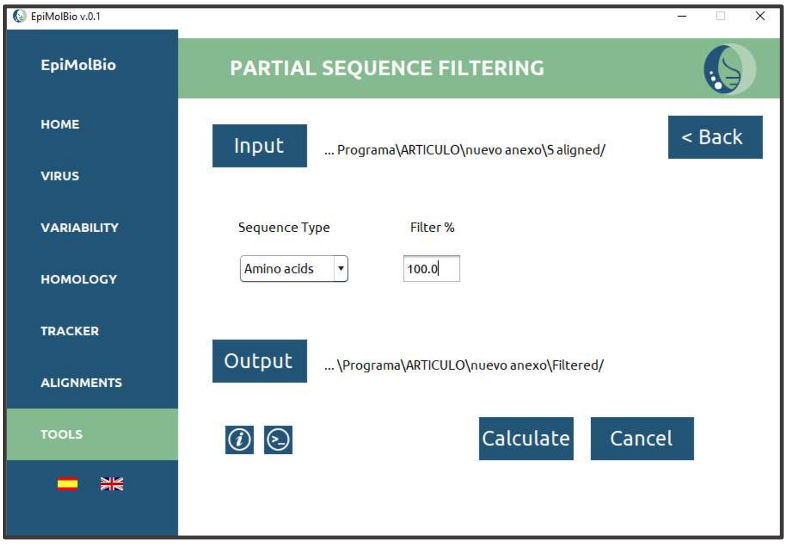


**Partial Sequence Filtering**

| File | Total Sequences | Recovered Sequences | Lost Sequences | Loss Percentage |
|---|---|---|---|---|
| Aligned_S (Spike)_Tracked_gisaid_hcov-19_2023_11_07_13.fas | 64 | 41 | 23 | 35.938% |
| Total | 64 | 41 | 23 | 35.938% |

The last step was to find the conserved sequence fragments in the Spike protein subunit 1 using the 'Search for conserved sequences' in EpiMolBio 'Homology' function.

The input was the previous output folder with the .fasta sequences. We wanted to search for these conserved fragments in the S1 of the Spike protein, which is between amino acids 14-685. Therefore, we chose 'Select Range' in the 'Range'box and typed 14 and 685 in the 'Select Range' boxes. In this way, the program searched within this specific region. Next, we typed 100.0 in the 'Conservation %' box, as we wanted these fragments to be completely conserved. In the 'Sequence Length' box, we typed the length of the desired fragment, in this case we chose between 20 and 25 residues. We input the reference sequence without spaces or line breaks. Finally, we created a new output folder named 'Conserved S' and named the file 'S1.html'.

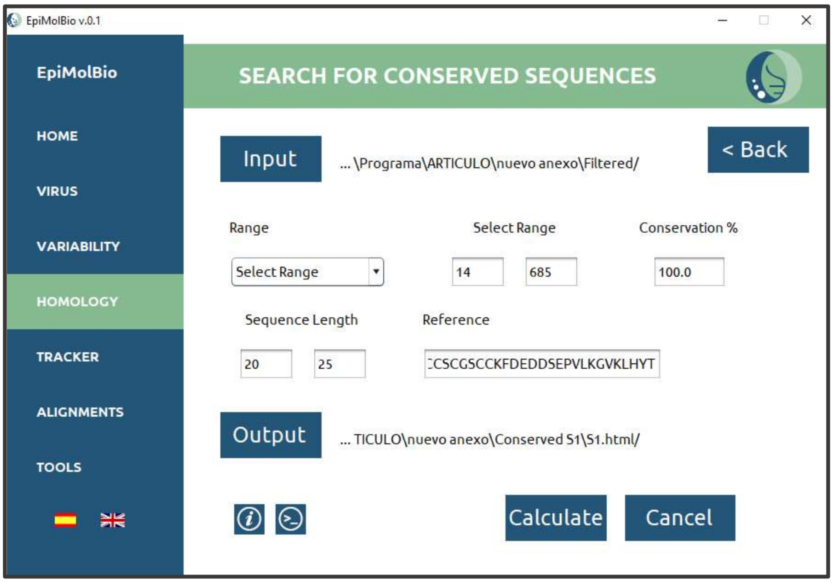


The resulting .html file showed a total of 756 completely conserved fragments within the SARS-CoV-2 Spike S1 with a length between 20-25 aa.

We show a partial result of the described analysis:

**Homology Search for Conserved Sequences Length 20 - 25 100.0%**

| File | Length | Region | Fragment | Frequency |
|---|---|---|---|---|
| Partial_Filter_Aligned_S (Spike)_Tracked_gisaid_hcov-19_2023_11_07_13.fas | 20 | 28 - 47 | YTNSFTRGVYYPDKVFRSSV | 100.000% |
| Partial_Filter_Aligned_S (Spike)_Tracked_gisaid_hcov-19_2023_11_07_13.fas | 20 | 29 - 48 | TNSFTRGVYYPDKVFRSSVL | 100.000% |
| Partial_Filter_Aligned_S (Spike)_Tracked_gisaid_hcov-19_2023_11_07_13.fas | 20 | 30 - 49 | NSFTRGVYYPDKVFRSSVLH | 100.000% |
| Partial_Filter_Aligned_S (Spike)_Tracked_gisaid_hcov-19_2023_11_07_13.fas | 20 | 84 - 103 | LPFNDGVYFASTEKSNIIRG | 100.000% |
| Partial_Filter_Aligned_S (Spike)_Tracked_gisaid_hcov-19_2023_11_07_13.fas | 20 | 85 - 104 | PFNDGVYFASTEKSNIIRGW | 100.000% |
| Partial_Filter_Aligned_S (Spike)_Tracked_gisaid_hcov-19_2023_11_07_13.fas | 20 | 86 - 105 | FNDGVYFASTEKSNIIRGWI | 100.000% |
| Partial_Filter_Aligned_S (Spike)_Tracked_gisaid_hcov-19_2023_11_07_13.fas | 20 | 87 - 106 | NDGVYFASTEKSNIIRGWIF | 100.000% |
| Partial_Filter_Aligned_S (Spike)_Tracked_gisaid_hcov-19_2023_11_07_13.fas | 20 | 88 - 107 | DGVYFASTEKSNIIRGWIFG | 100.000% |

In the 'File' column, the name of the analyzed file appears, followed by the 'Length' column displaying the chosen length of the fragment. The 'Region' column shows the region of the input sequence where the fragment has been found. In the 'Fragment' column, the result is displayed with the obtained sequence, followed by the 'Frequency' column, which shows the conservation percentage.

Shu, Yuelong, and McCauley, John, ‘GISAID: Global Initiative on Sharing All Influenza Data - from Vision to Reality.’, *Euro Surveillance : Bulletin Europeen Sur Les Maladies Transmissibles = European Communicable Disease Bulletin* (Sweden, 2017)

We gratefully acknowledge the Authors and their Originating laboratories responsible for obtaining the specimens, and their Submitting laboratories for generating the genetic sequence and metadata and sharing via the GISAID Initiative, on which this example is based.

**Data Availability**

GISAID Identifier: EPI_SET_231107fa
doi: 10.55876/gis8.231107fa

All genome sequences and associated metadata in this dataset are published in GISAID’s EpiCoV database. To view the contributors of each individual sequence with details such as accession number, Virus name, Collection date, Originating Lab and Submitting Lab and the list of Authors, visit 10.55876/gis8.231107fa

**Data Snapshot**

- EPI_SET_231107fa is composed of 65 individual genome sequences.
- The collection dates range from 2023-11-01 to 2023-11-02;
- Data were collected in 4 countries and territories;
- All sequences in this dataset are compared relative to hCoV-19/Wuhan/WIV04/2019 (WIV04), the official reference sequence employed by GISAID (EPI_ISL_402124). Learn more at https://gisaid.org/WIV04.
