## Supplementary Table 1 for "EpiMolBio: A Novel User-Friendly Bioinformatic Program for Genetic Variability Analysis"

**Supplementary Table 1. Fifty-three examples demonstrating various functions of EpiMolBio available on GitHub for training purposes** (https://github.com/EpiMolBio/EpiMolBio).

| Examples E1-E14 | | | | | Shown examples | Language | Example No. |
|---|---|---|---|---|---|---|---|
| VIRUS | HIV | Resistance Mutations | Individual | Acquired | 3 | SP | E1-E3 |
| | | | | | 3 | EN | E1-E3 |
| | | | | Transmitted | 3 | SP | E4-E6 |
| | | | | | 3 | EN | E4-E6 |
| | | | Codons | | 2 | SP | E7-E8 |
| | | | | | 2 | EN | E7-E8 |
| | | Pol Conservation | | | 1 | SP | E9 |
| | | | | | 1 | EN | E9 |
| | | Other Pol Mutations | | | 4 | SP | E10-E13 |
| | | | | | 4 | EN | E10-E13 |
| | SARS-CoV 2 | Protein Tracker | | | 1 | SP | E14 |
| | | | | | 1 | EN | E14 |
| | | | | TOTAL EXAMPLES | 28 | | E1-E14 |

| Examples E15-E33 | | | Shown examples | Language | Example No. |
|---|---|---|---|---|---|
| VARIABILITY | Polymorphisms | Individual | 9 | SP | E15-E23 |
| | | | 9 | EN | E15-E23 |
| | | Codons | 2 | SP | E24-E25 |
| | | | 2 | EN | E24-E25 |
| | Conservation | Individual | 2 | SP | E26-E27 |
| | | | 2 | EN | E26-E27 |
| | | Codons | 2 | SP | E28-E29 |
| | | | 2 | EN | E28-E29 |
| | Consensus | | 2 | SP | E30-E31 |
| | | | 2 | EN | E30-E31 |
| | Wu-Kabat | | 1 | SP | E32 |
| | | | 1 | EN | E32 |
| | Mutation Frequency | | 1 | SP | E33 |
| | | | 1 | EN | E33 |
| | | TOTAL EXAMPLES | 38 | | E15-E33 |

| Examples E34-E36 | | Shown examples | Language | Example No. |
|---|---|---|---|---|
| HOMOLOGY | Similarity | 1 | SP | E34 |
| | | 1 | EN | E34 |
| | Partial Similarity | 1 | SP | E35 |
| | | 1 | EN | E35 |
| | Search for Conserved Sequences | 1 | SP | E36 |
| | | 1 | EN | E36 |
| TOTAL EXAMPLES | | 6 | | E34-E36 |

| Examples E37-E38 | | Shown examples | Language | Example No. |
|---|---|---|---|---|
| TRACKER | Similarity | 1 | SP | E37 |
| | | 1 | EN | E37 |
| | Flanking | 1 | SP | E38 |
| | | 1 | EN | E38 |
| TOTAL EXAMPLES | | 4 | | E37-E38 |

| Examples E39-E41 | | Shown examples | Language | Example No. |
|---|---|---|---|---|
| ALIGNMENTS | Multiples Alignments | 1 | SP | E39 |
| | | 1 | EN | E39 |
| | Dot Plot | 1 | SP | E40 |
| | | 1 | EN | E40 |
| | Delete Insertions | 1 | SP | E41 |
| | | 1 | EN | E41 |
| TOTAL EXAMPLES | | 6 | | E39-E41 |

| Examples E42-E53 | | | Shown examples | Language | Example No. |
|---|---|---|---|---|---|
| TOOLS | File Editing | Merge Files | 1 | SP | E42 |
| | | | 1 | EN | E42 |
| | | Unique Sequences | 1 | SP | E43 |
| | | | 1 | EN | E43 |
| | | Sequence Search | 2 | SP | E44-E45 |
| | | | 2 | EN | E44-E45 |
| | | Find and Replace | 1 | SP | E46 |
| | | | 1 | EN | E46 |
| | Filters | Header Filtering | 1 | SP | E47 |
| | | | 1 | EN | E47 |
| | | Specific Filter | 1 | SP | E48 |
| | | | 1 | EN | E48 |
| | | Partial Sequence Filtering | 1 | SP | E49 |
| | | | 1 | EN | E49 |
| | Translation | | 1 | SP | E50 |
| | | | 1 | EN | E50 |
| | Count Sequences | | 2 | SP | E51-E52 |
| | | | 2 | EN | E51-E52 |
| | Function Programming | | 1 | SP | E53 |
| | | | 1 | EN | E53 |
| TOTAL EXAMPLES | | | 24 | | E42-E53 |

E, example; SP, in Spanish; EN, in English. Numbers refers to different provided examples.
