## Supplementary Table 2 for "EpiMolBio: A Novel User-Friendly Bioinformatic Program for Genetic Variability Analysis"

### Supplementary Table 2. Summary of EpiMolBio Program Functions

| Main Section | Sub-section | Function |
|---|---|---|
| **VIRUS** | HIV | |
| | Resistance Mutationsǂ (Acquired* and Transmitted) | Calculates the percentage of acquired/transmitted DRM, relative to a reference sequence, from amino acid (aa) sequences of HIV Pol proteins and p24 (Capsid). |
| | Other Pol Mutations | Detects any mutation (not only DRM) in HIV-1 or HIV-2 from sequences of Pol proteins, and their percentage relative to the reference sequence. |
| | Pol Conservation | Generates a table displaying the most prevalent aa and its corresponding percentage at each position within the selected Pol protein sequence, facilitating the identification of the protein's most conserved residue for each position |
| | SARS-CoV-2 Protein Tracker | Provides sequences in .fasta format of the chosen proteins from SARS-CoV-2, based on complete genomes, in nucleotides (nt) or aa. |
| **VARIABILITY** | Polymorphisms* | **Mutated Positions** and **Mutations Table** enable the detection of polymorphisms, providing information about their location and frequency of occurrence using any sequence introduced by the user as a reference.<br>**Markers** allows the detection of mutations exclusive to each file compared to the rest of the input files.<br>**Multiple Mutations** enables the detection of mutation combinations providing their frequency of occurrence.<br>**Codon function** detects the codons that are different from those in the reference sequence and their frequency of occurrence.<br>**Mutations by Position** allows the detection of residues at one position or several combined positions providing their frequency of occurrence. |
| | Conservation* | Determines the level of conservation of sequences of interest by reporting the most prevalent residue and its corresponding percentage. Additionally, it generates consensus sequences<br>**Codon function** presents the most conserved codon from an analyzed nt sequence. |
| | Consensus | Provides consensus sequences and consensus of consensuses by performing multiple rounds of analysis. |
| | Wu-Kabat Coefficient | Provides the Wu-Kabat variability coefficient of protein sequences to study the susceptibility of an aa position to evolutionary replacements. |
| | Mutation Frequency | Generates a set of parameters related to the frequency of mutations in a group of sequences, such as mutation frequency, conservation percentage, and average mutations per sequence. |
| **HOMOLOGY** | Similarity | Searchs for a user-introduced target sequence among the sequences in the input file, obtaining the proportion of sequences per file that contain the target sequence. |
| | Partial Similarity | Compares a user-introduced sequence with the input sequences to search for similar regions between them, defining the percentage of similarity between the sequences. |
| | Search for Conserved Sequences | Extracts conserved sequence fragments from a set of input sequences. Allows searching within a specific region, choosing the fragment length, and establishing the conservation percentage. |
| **TRACKER** | Similarity | Searchs for target sequences of interest within a set of longer sequences based on a reference sequence. |
| | Flanking | Searchs for proteins within a set of complete genomic sequences using the flanking sequences of the target protein. |
| **ALIGNMENTS** | Multiple Alignments | Aligns aa and nt sequences using the MUSCLE v3.8.31 program. |
| | Dot Plot | Generates a graphic where sequences are compared by plotting points on a two-dimensional matrix, with each axis representing a sequence. |
| | Delete Insertions | Automatically removes insertions from a sequence with respect to a reference with gaps after performing the alignment. |
| **TOOLS** | File Editing | |
| | Merge Files | Combines multiple .fasta files into a single .fasta file. |
| | Unique Sequences | Removes duplicated sequences from one or multiple input .fasta files. |
| | Sequence Search | Filters sequences from .fasta files that contain one or multiple mutations chosen by the user. |
| | Find and Replace | Replaces a series of characters with others in both the header and the genetic sequence of one or multiple .fasta files. |
| | Filters | |
| | Header Filtering | Filters one or multiple .fasta format files using parameters from their headers. |
| | Specific Filter | Filters sequences from .fasta format files that have a specific set of characters in their headers. |
| | Partial Sequence Filtering | Filters sequences from .fasta format files based on their quality, depending on the quantity of "?" (sequences in aa) or "N" (sequences in nt) they contain. |
| | Translation | Translates .fasta sequences from nt to aa. |
| | Count Sequences | Counts the total number of sequences in one or multiple .fasta files, or how many of those sequences contain mutations. |
| | Function Programming | Automates the program's functions by chaining them together to be executed sequentially without manual intervention. |

ǂ DRM according to Stanford HIV Drug Resistance Database v9.7 for HIV-1 (https://hivdb.stanford.edu/page/release-notes/#comments), and to HIV-2 EU Tool v.2 and recent literature for HIV-2 (see manuscript).
* Individual and codon analysis
Abbreviations: DRM, drug resistance mutations;aa, amino acid; nt, nucleotide
